## Supplementary material for "Illuminating the plant rhabdovirus landscape through metatranscriptomics data": Supp Material: Supp Table S3.docx

Supplementary Table S3. Characteristics of deduced proteins encoded by each assembled rhabdovirus sequence determined by predictive algorithms

| **Virus** | **RNA** | **ORF No^a^** | **Gene name** | **Mr (kDa)** | **IEP** | **TMHMM** | **Signal P cleavage site** | **Predicted NLS**  **[cNLS Mapper score] #** | **Predicted NES** | **Highest scoring virus- protein/*E*-value/query coverage% (Blast P)** |
| --- | --- | --- | --- | --- | --- | --- | --- | --- | --- | --- |
| *Agave tequilana* virus 1 | 1 | 1  2  3  4  5  6 | N  P  P3  M  G  L | 50  37.4  32  28.8  67.8  221.2 | 8.49  5.58  8.54  6.71  6.45  6.97 | -  -  -  -  551-573  - | -  -  -  -  19:20  - | ^429^TPKKKKRTGD^438^ [9]  ^289^TSKKRYESDPAFNKIGDEDLEKCKKLKSE^317^ [6.5]  ^197^DIIKKASMAANHLHLPEDKALISNPSNIIKL^227^ [3.4]  ^167^FGSVKEPSGDPQVLKTLIDIQFGCLLKNPSTT^198^ [4.2]  ^277^IYSIKRKISAMPLKDRIELLWKRTPET^303^ [4.6]  ^735^RETLKRIMEDLTCFLSKLGLPLKRAETW^762^ [7.7] | 276  313-324  -  -  -  1416, 1419 | MMaV-N/4e^71/86  no hits  PeV1-P3/0.012/56  MIMV-M/2e^04/61  TaVCV-G/5e-90/90  MMV-L/0.0/94 |
| *Allium angulosum* virus 1 | 1  2 | 1  1  2  3 | L  N  2  3 | 236.6  54.9  51.2  15.2 | 7.91  5.18  9.06  8.33 | -  -  -  - | -  -  -  - | ^1542^ YTREKKKAKLEYIEC^1556^ [9.5]  -  ^189^DQKKKRKLE^197^ [8.5]  - | 36, 234, 1933  -  392-394, 397  - | AMVV1-L/0.0/94  AMVV1-N/2e^41/68  no hits  no hits |
| *Allium chinense* virus 1 | 1 | 1  2  3  4  5  6  7  8 | N  P  P’  P3  M  G*  P6*  L* | 52.1  35.4  15.4226.1  18.5  -  -  - | 6.29  4.57  10.1  9.59  9.18  5.82  -  - | -  -  42-61, 82-107  -  -  492-514  -  - | -  -  -  -  -  20:21  -  - | -  -  -  -  -  -  -  - | 308  -  -  -  119-126  -  -  53, 55, 418, 420 | SCV-N/3e^140/96  ChYDaV-P/7e^60/100  RVCV-P’/0.001/34  ChYDaV-P3/3e^73/96  ChYDaV-M/6e^25/99  RVCV-G/6e^143/95  -  SCV-L/7e-70/100 |
| *Anthurium amnícola* virus 1 | 1 | 1  2  3  4  5  6 | N  P  P3  M  G  L | 45.6  35.9  23.7  20.7  63.8  234.2 | 5.82  5.11  7.75  9.54  8.15  8.48 | -  -  -  -  13-35, 40-59, 526-548  - | -  -  -  -  -  - | -  -  -  -  -  - | 369  -  -  36, 126  -  88-89, 91 1259,1344, 1349 | PMuMaV-N/1e^104/99  PMuMaV-P/2e^11/64  PMuMaV-P3/3e^23/76  PMuMaV-M/9e^33/98  PMuMaV-G/3e^76/96  PMuMaV-L/0.0/99 |
| *Asclepias syriaca* virus 1 | 1 | 1  2  3  4  5  6  7  8 | N  P  P’  P3  M  G  P6  L | 50  33.8  10.2  37.7  19.6  61.8  10.02  241.6 | 5.7  4.58  9.75  8.78  9.16  6.34  9.52  8.86 | -  -  5-27, 32-51, 64-86  -  -  502-524  27-49  - | -  -  -  -  -  22:23  -  - | -  -  -  -  -  -  -  - | 100, 102  -  -  -  128-134  -  -  - | WhIV4-N/1e^164/93  WhIV4-P/8e^48/89  no hits  WhIV4-P3/4e^123/99  WhIV-4-M/4e-23/91  WhIV-4-G/6e^161/99  no hits  WhIV4-L/0.0/99 |
| *Asclepias syriaca* virus 2 | 1 | 1  2  3  4  5  6 | N  P  P3  M  G  L | 51.8  38.2  39.3  28.3  73.5  233.4 | 8.73  4.86  8.62  5.95  6.43  7.23 | -  -  -  -  560-582  - | -  -  -  -  25:26  - | ^413^IVARKRGHTLGTIPDKPNTATKRSASE^439^ [5.2]  ^82^LHKNNFNQLSDVLLDGIVTFGKEIRLSTVP^111^ [4.1]  ^70^ESYLNKRKIELSLKNLFNVFSVEVDIKQITCIW^102^ [4.8]  ^147^LDKDHGVIVSNIEIFISPLTAEQFKASKL^175^ [5.5]  ^250^RIDCHYKESSAILECPEITYSFPLHRLKKTPTCK^283^ [4.6]  ^790^EKMQKRGLPLKLEETWISNRLLMYNKIMYL^819^ [5.8] | 86  -  -  -  -  - | ApRVA-N/5e^146/98  ApRVA-P/3e^11/69  ApRVA-P3/4e^36/95  ApRVA-M/53^18/82  ApRVA-G/5e^133/95  ApRVA-L/0.0/99 |
| *Bemisia tabaci* -associated virus 1 | 1 | 1  2  3  4  5  6  7 | N  P  P3  P4  M  G  L | 50.2  36.1  21.6  5.99  23.8  65.7  242.1 | 8.67  4.64  9.35  10.29  8.96  7.46  8.43 | -  -  -  5-22  -  22-39, 59-78, 538-557  - | -  -  -  -  -  - | -  -  -  -  -  -  - | 61  275  -  10-16  -  -  573, 577, 1423, 1465, 1983, 1985-1986, 1988, 1990 | CuCV1-N/0.0/100  CuCV1-P/1e^148/96  CuCV1-P3/3e^96/100  CuCV1-P4/6e^06/72  CuCV1-M/2e^93/100  CuCV1-G/0.0/100  CuCV1-L/0.0/99 |
| *Brassica rapa* virus 1 | 1  2 | 1  1  2  3 | L  N  2  3 | 232.4  49.9  46.3  20.7 | 6.92  5.29  9.18  5.03 | -  -  -  - | -  -  -  - | _1594_FFKMNPSLIEDYGEEEDFKIKAEIKQFRRK^1623^ [5.6] ^221^KLVERFEHFYMANFPFEDFHPNLESAKAIKSI^252^ [5.4]  ^287^EPRLKRKESPSPPILDVKKPKKSESK^312^ [6.8]  - | 6, 11, 2004  373, 375, 378, 380  70, 190  147 | RCaVV-L/0.0/99  RCaVV-N/1e^43/72  no hits  RCaVV-P3/2e-09/90 |
| *Cuscuta reflexa* virus 1 | 1 | 1  2  3  4  5  6 | N  P  P3  M*  G*  L* | 50.8  36.8  36.8  -  -  - | 6.72  5.51  8.48  8.89  5.55  - | -  -  -  -  479-501  - | -  -  -  -  23:24  - | ^440^EPTKKRKTPSL^450^ [8]  ^218^EQLRAKVRLHYSDTEFDSFNDSKKMAL^244^ [4]  ^114^RGRISESLKAVVFPTKSILKNKNDAFFMPWKFAA^147^ [4]  ^200^RQKRVPRKRSKSPAYNTKKRGISKKRKPQ^228^ [8.8]  ^507^KVPSKVVTFVEDEMEDFKTPLRVPSAPKKRM^537^ [5]  ^288^EGVGQKRQHPSVQESIKRHMLSDRTHATRMRLS^320^ [5.3] | 292  214-225  171, 173  -  -  - | DYVV-N/0.0/97  DYVV-P/8e^53/97  DYVV-P3/1e^91/91  GSPNuV-M/2e^77/100  DYVV-G/0.0/92  DYVV-L/3e^71/98 |
| *Dioscorea composita* virus 1 | 1 | 1  2  3  4  5  6  7 | N  P*  P3*  M  G  P6  L* | 48.5  -  -  22.9  66.2  8.54  - | 6.37  -  -  7.19  6.51  11.18  - | -  -  -  -  539-556  23-45  - | -  -  -  -  -  -  - | -  -  -  -  -  -  - | 148  -  -  13  -  -  37 | MYSV-N/5e^23/93  -  -  no hits  PMuMaV-G/4e^04/85  no hits  RVR/0.0/95 |
| *Glehnia littoralis* virus 1 | 1 | 1  2  3  4  5  6  7  8 | N  P  P’  P3  M  G  P6  L | 46.4  35.2  10.3  22.6  18.6  60.9  7.8  238.3 | 5.6  5.47  10.4  9.41  8.48  6.5  9.74  8.23 | -  -  13-35, 50-72  -  -  503-525  31-53  - | -  -  -  -  -  21:22  -  - | -  -  -  -  -  -  -  - | 264  -  -  -  -  -  -  95, 97-98, 100 | TpVA-N/0.0/99  TpVA-P/6e^136/99  no hits  TpVA-P3/3e^114/100  TpVA-M/8e^79/94  TpVA-G/0.0/98  no hits  TpVA-L/0.0/99 |
| *Gymnadenia densiflora* virus 1 | 1 | 1  2  3  4 | N  P  M  L | 49  34.6  21.7  237.7 | 5.5  6.49  5.5  7.81 | -  -  -  - | -  -  -  - | -  -  -  - | 73, 80-86  124  155  1574 | LYMoV-N/6e^33/89  SCV-P/0.002/42  no hits  WhIV5-L/0.0/88 |
| *Lolium perenne* virus 1 | 1  2 | 1  1  2  3 | L  N  2  3 | 231.9  61.9  43  18.3 | 7.19  6.48  5.84  5.34 | -  -  -  - | -  -  -  - | ^108^RLVHKALKNRDIRIPISTNDITEDLPDVKIYHRW^141^ [5.4]  ^134^VAPKKKKKRMRHF^146^ [8.5]  ^91^GKKKKPRLGRE^101^ [8]  - | -  -  163, 165  93-98 | AMVV1-L/0.0/100  AMVV1-N/2e^142/94  no hits  no hits |
| *Lotus corniculatus* virus 1 | 1 | 1  2  3  4  5  6  7  8 | N  P  P’  P3  M  G  P6  L* | 53.7  34.6  11.8  40.2  19.8  62.6  6.65  - | 6.36  4.71  10.6  8.56  8.75  6.97  9.57  8.31 | -  -  30-47, 51-73  -  -  512-534  21-43  - | -  -  -  -  -  24:25  -  - | -  -  -  -  -  -  -  - | 128-130  252, 262  -  -  123-125  -  -  832-833, 901, 905, 907, 909, 1484,1915, 1920, 1922 | LNYV-N/9e^77/96  WhIV4-P/2e^25/57  no hits  WhIV4-P3/2e^50/74  CCyV1-M/0.001/53  WhIV4-G/1e^101/98  no hits  WhIV4-L/0.0/98 |
| *Melampyrum roseum* virus 1 | 1  2 | 1  1  2  3  4 | L  N  2  3  4 | 229.2  49.8  38.1  34.5  22 | 7.35  6.04  5.34  9.1  5.42 | -  -  -  -  - | -  -  -  -  - | ^107^IGDALKKRKID^117^ [9.5]  -  -  - | -  23  37  71-78  81 | VVV-L/0.0/96  VVV-N/8e^71/91  no hits  VVV/2e^74/87  no hits |
| *Nymphaea alba* virus 1 | 1 | 1  2  3  4  5  6  7  8 | N  P  P’  P3  M  G  P6  L | 47.4  33.3  10  25.4  21.3  62.5  8.3  237.7 | 5.8  4.8  9.1  8.68  8.94  6.48  9.78  8.46 | -  -  7-29, 33-52, 57-79  -  -  504-526  32-54  - | -  -  -  -  -  22:23  -  - | -  -  -  -  -  -  -  - | -  287  -  -  36  -  -  - | TpVA-N/6e^115/95  TpVA-P/1e-22/99  no hits  StrV1-P3/3e^32/69  KePCyV-M/3e^11/50  KePCyV-G/1e^146/93  no hits  TpVA-L/0.0/98 |
| *Pelargonium radula* virus 1 | 1 | 1  2  3  4  5  6  7  8 | N  P  P’  P3  M  G*  P6  L* | 53.9  33.9  8.57  39.1  19.5  -  6.66  - | 7.07  4.51  10.1  8.63  8.3  6.04  10.2  - | -  -  4-26, 33-55  -  -  20-42, 529-551  15-37  - | -  -  -  -  -  41:42  -  - | -  -  -  -  -  -  -  - | 135  277, 281  -  -  61  -  -  89, 91,112 | WhIV4-N/3e^121/98  WhIV4-P/4e-27/90  no hits  WhIV4-P3/5e^73/94  WhIV4-M/2e^06/85  WhIV4-G/1e^86/93  no hits  WhIV4-L/0.0/97 |
| *Persicaria minor* virus 1 | 1 | 1  2  3  4  5  6 | N*  P*  P3*  M*  G*  L* | -  -  -  -  -  - | -  -  -  -  -  - | -  -  -  -  -  - | -  -  -  -  -  - | ^47^LFKIPGALNNCALIGPPTSHQRAAKCQWCK^76^ [4.9]  ^51^NLFKAEGLTIPNETISVLGEVSLVCKKHGRE^80^ [3.5]  -  ^183^PEKKKRPRTS^192^ [8]  ^202^RNPGWFSKVGLTRSMLHSQADIAGISSPRCRPIKK^236^ [3.7]  ^356^TAEKFKTGVLLSAYGDEDLLKRALKIQKL^384^ [7] | -  -  -  120-122, 124  -  66, 73, 303-304 | GSPNuV-N/7e^131/97  DYVV-P/4e^18/72  DYVV-P3/3e^42/81  GSPNuV-M/2e^63/96  GSPNuV-G/6e^72/95  DYVV-L/4e^45/67 |
| *Phlox pilosa* virus 1 | 1  2 | 1  2  1  2  3  4  5 | 6*  L*  N  2*  3  4*  5* | -  -  42.3  -  31.7  -  - | -  -  5.9  -  8.74  -  - | -  -  -  -  -  -  - | -  -  -  -  -  -  - | -  -  -  -  -  -  - | -  79, 81, 83  67, 69  -  236, 239-242, 244  25-29  - | -  LBVaV-L/1e^154/97  LBVaV-N/3e^45/99  no hits  LBVaV-P3/3e^50/97  no hits  no hits |
| *Pinus flexilis* virus 1 | 1 | 1  2  3  4  5 | N  2  3  4  L | 45.7  50.2  35.4  25.2  232.6 | 8.28  4.92  8.8  5.32  7.54 | -  -  -  -  - | -  -  -  -  - | -  ^79^KVLKRVKDEGEGSRQSPKDKGKTRKNPFVI^108^ [5.9]  ^232^PRRPSSKMIQTYARRGVNSASRATKIIKL^250^ [5.9]  -  - | 321  191, 193, 196, 198  278, 280  168  26 | AMVV1-N/4e^18/86  no hits  LBVaV-P3/5e^37/80  no hits  LBVaV-L/0.0/97 |
| *Plectranthus aromaticus* virus 1 | 1 | 1  2  3  4  5  6 | N  P  P3  M  G  L | 50.2  36.7  36.1  32  65.9  238.1 | 8.13  5.16  8.03  8.71  5.91  8.26 | -  -  -  -  500-522  - | -  -  -  -  -  - | ^421^DNEAKRKAPISEDETAKKKRPPP^443^ [11.5]  ^140^RKKVSIAKKIEGKDITLPPSQESQSKDPGDKTIK^173^ [5.1]  ^178^ERVLYKPNVVSWSNLHYPYYIPFYMEKKVRGI^209^ [3.4]  ^220^RPKQRFNKRPGLSPSSAPAKKILN^243^ [5.5]  ^538^REKIVKFNPDALEQFVPDNLVHPTAPKRRPQS^569^ [4.1]  ^1619^DQTTSKRKRAGGLFSYIKRARD^1640^ [9.8] | 350  73  -  150, 154  -  166, 1455 | DYVV/0.0/97  DYVV-P/8e^93/97  DYVV-P3/1e^160/100  DYVV-M/8e^97/94  DYVV-G/0.0/100  DYVV-L/0.0/100 |
| *Rhododendron delavayi* virus 1 | 1 | 1  2  3  4  5  6 | N  P  P3  M  G  L | 51.1  38.3  36.9  31.2  73  233 | 8.6  5.19  8.16  5.77  8.54  8.37 | -  -  -  -  558-577  - | -  -  -  -  23:24  - | ^436^RGKRPLEQEGGIPKRPAMESVPVAPLPFST^465^ [4.6]  ^227^RDRCRKHYTSELFESWDDAKKRTSI^251^ [8.8]  ^230^DHCKRLKISEEDSWNLMQTMTHEDANKIMDKT^261^ [4.1]  ^225^KSYLKHLLSRVGKRGRDDSPYQLRKEKLSKT^255^ [7.4]  ^82^KQACLNEQKETEITIAIMKWDFKTDKIP^109^ [5.4]  ^1656^KKGKRKIGSLQELLNRGAAAKRIRVS^1681^ [9] | -  -  236  -  -  1458 | SYVV-N/9e^126/89  BCaRV-P/7e^30/97  BCaRV-P3/2e^59/91  BCaRV-M/6e^29/79  DYVV-G/1e^132/96  BCaRV-L/0.0/95 |
| Spinach virus 1 | 1  2 | 1  1  2  3 | L*  N  2  3* | 231.3  50.6  50.5  - | 7.34  6.15  5.59  - | -  -  -  - | -  -  -  - | ^1936^KATKRNRLD^1944^ [5.5]  -  ^5^AIKRTSQATLQEGEDTRKSTRL^26^ [5]  - | -  292  377, 382  63 | RCaVV-L/0.0/99  RCaVV/4e^42/91  no hits  RCaVV-P3/8e^04/90 |
| *Suaeda salsa* virus 1 | 1 | 1  2  3  4  5  6  7  8 | N*  P*  P’*  P3  M  G*  P6*  L* | -  -  -  24.4  18.2  -  -  - | -  4.72  -  9.41  9.52  -  -  - | -  -  36-58  -  -  -  -  - | -  -  -  -  -  -  -  - | -  -  -  -  -  -  -  - | 172, 175, 178  -  -  -  119-126  -  294 | SCV-N/6e^107/97  RVCV-P/4e^46/82  no hits  ChYDaV-P3/1e^55/85  RVCV-M/8e^25/96  ChYDaV-G/2e^14/98  no hits  RVCV-L/8e^119/80 |
| *Tagetes erecta* virus 1 | 1 | 1  2  3  4  5 | N  P  P3  M  L | 59.9  56.6  23.1  21.1  238.8 | 5.47  5.48  9.57  8.84  7.04 | -  -  -  -  - | -  -  -  -  - | -  -  -  -  - | 69, 72  -  -  -  123, 1957, 1959,1961 | RSMV-N/8e^42/66  PpVE-P/1e^09/55  RVR-P3/3e^20/66  MYSV-M/0.001/45  MYSV-L/0.0/99 |
| *Trachyspermum ammi* virus 1 | 1 | 1  2  3  4  5 | N  P  P3  M  L | 50.4  36.6  26.5  22.8  236.8 | 6.02  7.64  8.43  4.71  7.26 | -  -  -  -  - | -  -  -  -  - | -  -  -  -  - | 88-92  131  -  60, 62-63  - | StrV1-N/4e^30/62  no hits  StrV1-P3/0.002/49  CCyV1-M/2e^04/44  WhIV5-L/0.0/96 |
| *Viola verecunda* virus 1 | 1  2 | 1  2  3  4  5  1 | N  P*  P3  M*  G  L* | 55.8  -  36.6  -  74.4  - | 5.3  -  8.59  -  5.54  - | -  -  -  -  618-640  - | -  -  -  -  -  - | ^427^PRPEKRKRADRE^438^ [10]  -  ^47^RDVVCKGKFGPNLLRRWGNSTHHSVCVKEIKID^79^ [5.6]  -  ^31^KQTPEQAPADPDVRPERELKDEKMTEM^57^ [3.6]  ^1655^IHPKGKKRKMK^1665^ [11.5] | 26, 176  -  -  -  -  1466 | OFV-N/2e^35/72  -  ORF-P3/7e^05/61  -  OFV-G/4e^05/47  ClCSV-L/1e^153/60 |

^a^ ORF numbers are represented from 3´ to 5´ for genomic sense and correspond to those shown in Fig.1.

* Coding region not complete

\# The predicted NLS with the highest score is shown

TMHMM=Transmembrane domain.

NLS=Nuclear localization signal

NES=Leucine-rich nuclear export signalNames and abbreviations of viruses are listed in Supp. Table S1.
