## Supplementary material for "Illuminating the plant rhabdovirus landscape through metatranscriptomics data": Supp Material: Supp.Table S1.docx

Supplementary Table S1. Virus names, abbreviations and NCBI accession numbers of plant rhabdovirus sequences used in this study

| **Virus name** | **Abbreviation** | **Accession number** |
| --- | --- | --- |
| actinidia cytorhabdovirus | AcCV | MW550041 |
| alfalfa-associated nucleorhabdovirus | AaNV | MG948563 |
| alfalfa dwarf virus | ADV | KP205452 |
| Alopecurus myosuroides varicosavirus 1 | AMVV1 | LN713933; LN713934 |
| apple rootstock virus A | ApRVA | MH778545 |
| Bacopa monnieri virus 1 | BmV1 | BK014479 |
| Bacopa monnieri virus 2 | BmV2 | BK014480 |
| barley yellow striate mosaic virus | BYSMV | KM213865 |
| bean associated rhabdovirus | BaCV | MK202584 |
| bird’s-foot trefoil virus 1 | BFTaV | BK010826 |
| black currant-associated rhabdovirus | BCaRV | MF543022 |
| cabbage cytorhabdovirus 1 | CCyV1 | KY810772 |
| cardamom vein clearing virus 1 | CdVCV1 | MN273311 |
| citrus-associated rhabdovirus | CiaRV | MT302547 |
| citrus chlorotic spot virus | CiCSV | KY700685; KY700686 |
| citrus leprosis virus N | CiLV-N | KX982176; KX982179 |
| clerodendrum chlorotic spot virus | ClSCV | MG938506: MG938507 |
| chrysanthemum yellow dwarf-associated virus | ChYDaV | MW039593 |
| coffee ringspot virus | CoRSV | KF812525; KF812526 |
| Colocasia bobone disease-associated virus | CBDaV | KT381973 |
| constricta yellow dwarf virus | CYDV | KY549567 |
| cucurbit cytorhabdovirus 1 | CuCV1 | MT381995 |
| datura yellow vein virus | DYVV | KM823531 |
| eggplant mottle dwarf virus | EMDV | KJ082087 |
| green Sichuan pepper nucleorhabdovirus | GSPNuV | MH323437 |
| Iranian citrus ringspot-associated virus | IrCRSaV | KP255975 |
| joa yellow blotch-associated virus | JYBaV | MW014292 |
| Kenyan potato cytorhabdovirus | KePCyV | MN689395 |
| lettuce big-vein associated virus | LBVaV | AB075039; AB114138 |
| lettuce necrotic yellows virus | LNYV | AJ867584 |
| lettuce yellow mottle virus | LYMoV | EF687738 |
| maize associated cytorhabdovirus | MaCyV | KY965147 |
| maize fine streak virus | MFSV | AY618417 |
| maize Iranian mosaic virus | MIMV | MF102281 |
| maize mosaic virus | MMV | MK828539 |
| Morogoro maize-associated virus | MMaV | MK112501 |
| maize yellow striate virus | MYSV | KY884303 |
| northern cereal mosaic virus | NCMV | MH282832 |
| orchid fleck virus | OFV | LC222629; LC222630 |
| papaya virus E | PpVE | MK202584 |
| paper mulberry mosaic-associated virus | PMuMaV | MN872813 |
| peach virus 1 | PeV1 | MN520414 |
| persimmon virus A | PeVA | AB735628 |
| Physostegia chlorotic mottle virus | PhCMoV | KY859866 |
| potato yellow dwarf virus | PYDV | GU734660 |
| raspberry vein chlorosis virus | RVCV | MK257717 |
| red clover-associated varicosavirus | RCaVV | MF918568; MF918569 |
| rice stripe mosaic virus | RSMV | MH720469 |
| rice yellow stunt virus | RYSV | AB011257 |
| rose virus R | RVR | MT952336 |
| sonchus yellow net virus | SYNV | L32603 |
| sowthistle yellow vein virus | SYVV | MT185675 |
| strawberry crinkle virus | SCV | MH129615 |
| strawberry virus 1 | StrV1 | MK211270 |
| taro vein chlorosis virus | TaVCV | AY674964 |
| tomato yellow mottle-associated virus | TYMaV | KY075646 |
| Trichosanthes-associated rhabdovirus 1 | TrARV1 | BK011194 |
| Trifolium pratense virus A | TpVA | MH982250 |
| Trifolium pratense virus B | TpVB | MH982249 |
| vitis varicosavirus | VVV | LC604719; LC604720 |
| wheat yellow striate virus | WYSV | MG604920 |
| Wuhan insect virus 4 | WhIV4 | KM817650 |
| Wuhan insect virus 5 | WhIV5 | KM817651 |
| Wuhan insect virus 6 | WhIV6 | KM817652 |
| yerba mate chlorosis-associated virus | YmCaV | KY366322 |
| yerba mate virus A | YmVA | MN781667 |
