## Supplementary material for "Illuminating the plant rhabdovirus landscape through metatranscriptomics data": Supp Material: Table S2.docx

**Table S2**. Summary of assembly statistics of the plant rhabdoviruses sequences identified from the transcriptome data available in the NCBI database.

| **Virus name** | **Abbreviation** | **Bioproject ID** | **SRA Accesion** | **Total virus reads** | **Mean Coverage** | **Reads per Millon** |
| --- | --- | --- | --- | --- | --- | --- |
| *Agave tequilana* virus 1 | ATV1 | PRJNA193469 | SRR789743 | **26874** | **306.54X** | **297.20** |
| *Asclepias syriaca* virus 2 | AscSyV2 | PRJNA210776 | SRR5117431 | **21120** | **164.84X** | **439.21** |
| *Cuscuta reflexa* virus 1 | CusReV1 | PRJNA290291 | SRR2142348 | **119** | **6.88X** | **63.99** |
| *Persicaria minor* virus 1 | PerMiV1 | PRJNA208436 | SRR917962 | **885** | **97.12X** | **1472.05** |
| *Plectranthus aromaticus* virus 1 | PleArV1 | PRJNA491230 | SRR7896533 | **17676** | **156.43X** | **2813.02** |
| *Rhododendron delavayi* virus 1 | RhoDeV1 | PRJNA358123 | SRR5121284 | **7841** | **51.43X** | **98.91** |
| *Allium chinense* virus 1 | AChV1 | PRJNA310810 | SRR3144560 | **878** | **13.75X** | **7.86** |
| *Anthurium amnícola* virus 1 | AntAmV1 | PRJNA288827 | SRR2089239 | **51637** | **620.63X** | **1495.45** |
| *Asclepias syriaca* virus 1 | AscSyV1 | PRJNA210776 | SRR5117431 | **51339** | **387.18X** | **1067.65** |
| *Bemisia tabaci* -associated virus 1 | BeTaV1 | PRJNA237273 | SRR1159208 | **88609** | **687.10X** | **1057.02** |
| *Dioscorea composita* virus 1 | DiCoV1 | PRJNA253902 | SRP044768 | **19538** | **196.18X** | **40.64** |
| *Glehnia littoralis* virus 1 | GlLV1 | PRJNA248158 | SRP042106 | **11812** | **97.84X** | **110.35** |
| *Gymnadenia densiflora* virus 1 | GymDenV1 | PRJNA504609 | SRR8175725 | **29407** | **301.22X** | **593.66** |
| *Lotus corniculatus* virus 1 | LotCorV1 | PRJNA77207 | SRR364671 | **3535** | **25.06X** | **132.41** |
| *Nymphaea alba* virus 1 | NymAV1 | PRJNA472003 | SRR7224571 | **135173** | **1048.99X** | **398.62** |
| *Pelargonium radula* virus 1 | PelRaV1 | PRJNA491235 | SRR7900215 | **676** | **6.07X** | **46.94** |
| *Suaeda salsa* virus 1 | SuSV1 | PRJNA395283 | SRP113741 | **520** | **8.67X** | **1.33** |
| *Tagetes erecta* virus 1 | TaEV1 | PRJNA431782 | SRR6667681 | **14945** | **159.57X** | **344.56** |
| *Trachyspermum ammi* virus 1 | TrAV1 | PRJNA359623 | SRR5137053 | **56323** | **515.77X** | **988.19** |
| *Viola verecunda* virus 1 | VVeV1 | PRJNA345302 | SRR5322180 | **453** | **4.35X** | **68.10** |
| *Allium* *angulosum* virus 1 | AAnV1 | PRJNA542932 | SRR9077160 | **18132** | **129.06X** | **142.44** |
| *Brassica rapa* virus 1 | BrRV1 | PRJNA396268 | SRR5914717 | **22527** | **269.07X** | **598.70** |
| *Lolium perenne* virus 1 | LoPV1 | PRJNA222646 | SRP044151 | **16413** | **158.34X** | **152.89** |
| *Melampyrum roseum* virus 1 | MelRoV1 | PRJDB5395 | DRR082665 | **12274** | **100.42X** | **183.59** |
| *Phlox pilosa* virus 1 | PhPiV1 | PRJNA360978 | SRP096783 | **5140** | **79.08X** | **99.70** |
| *Pinus flexilis* virus 1 | PiFleV1 | PRJNA315892 | SRR10032949 | **3076** | **26.20X** | **40.25** |
| Spinach virus 1 | SpV1 | PRJDB3392 | DRR029417 | **4166** | **42.08X** | **85.98** |
